# A fluid biomarker reveals loss of TDP-43 splicing repression in pre-symptomatic ALS

**DOI:** 10.1101/2023.01.23.525202

**Authors:** Katherine E. Irwin, Pei Jasin, Kerstin E. Braunstein, Irika Sinha, Kyra D. Bowden, Abhay Moghekar, Esther S. Oh, Denitza Raitcheva, Dan Bartlett, James D. Berry, Bryan Traynor, Jonathan P. Ling, Philip C. Wong

## Abstract

Loss of TAR DNA-binding protein 43 kDa (TDP-43) splicing repression is well-documented in postmortem tissues of amyotrophic lateral sclerosis (ALS), yet whether this abnormality occurs during early-stage disease remains unresolved. Cryptic exon inclusion reflects functional loss of TDP-43, and thus detection of cryptic exon-encoded peptides in cerebrospinal fluid (CSF) could reveal the earliest stages of TDP-43 dysregulation in patients. Here, we use a newly characterized monoclonal antibody specific to a TDP-43-dependent cryptic epitope (encoded by the cryptic exon found in *HDGFL2*) to show that loss of TDP-43 splicing repression occurs in *C9ORF72*-associated ALS, including pre-symptomatic mutation carriers. In contrast to neurofilament light and heavy chain proteins, cryptic HDGFL2 accumulates in CSF at higher levels during early stages of disease. Our findings indicate that loss of TDP-43 splicing repression occurs early in disease progression, even pre-symptomatically, and that detection of HDGFL2’s cryptic neoepitope may serve as a prognostic test for ALS which should facilitate patient recruitment and measurement of target engagement in clinical trials.

## Introduction

A fluid biomarker for the pre-symptomatic or prodromal phases of amyotrophic lateral sclerosis (ALS) to enable earlier diagnosis and to facilitate patient recruitment and monitor target engagement in clinical trials is a great unmet need. A central pathological hallmark of the amyotrophic lateral sclerosis &#8211; frontotemporal dementia (ALS-FTD) disease spectrum is the nuclear mislocalization and cytoplasmic aggregation of an RNA-binding protein termed TAR DNA-binding protein 43 kDa (TDP-43) (Neumann et al., 2006). While a gain-of-function mechanism due to TDP-43 cytoplasmic aggregates has been proposed to contribute to neurodegeneration (Barmada et al., 2010; Brettschneider et al., 2013; Lee et al., 2011; Taylor et al., 2016; Zhang et al., 2009), emerging evidence supports the idea that loss of TDP-43 splicing repression resulting from depletion of nuclear TDP-43 drives neuron loss in ALS-FTD (Ling et al., 2015; Tan et al., 2016). TDP-43 pathology can currently only be revealed postmortem, so while such TDP-43 functional deficits are well-documented in end-stage tissues (Brown et al., 2022; Klim et al., 2019; Ma et al., 2022; Melamed et al., 2019; Prudencio et al., 2020), the extent to which loss of TDP-43 splicing repression occurs during the early stages of disease is unclear. Clarifying this question would provide critical insight into disease mechanisms and inform therapeutic strategies designed to attenuate neuron loss in ALS-FTD.

Loss of TDP-43 splicing repression leads to the inclusion of numerous nonconserved cryptic exons, of which ~3% produce in-frame neoepitopes (Jeong et al., 2017). We hypothesize that detecting peptides encoded by cryptic exons in biofluids could reveal how early TDP-43 splicing repression is dysregulated in patients with ALS or FTD and could establish fluid biomarkers that reflect TDP-43 dysfunction (Fig. S1). To test this idea, we selected certain cryptic neoepitopes for antibody generation based on RNA expression data and protein structure modeling. We then validated these novel monoclonal antibodies in HeLa cells depleted of TDP-43 by siRNA. We focus here on one antibody that was able to reliably detect a cryptic exon-encoded neoepitope in Hepatoma-Derived Growth Factor-Like protein 2 (HDGFL2). Using this novel antibody, we developed a highly specific and sensitive sandwich enzyme-linked immunosorbent assay (ELISA) to determine the dynamic nature of this cryptic peptide target in CSF from pre-symptomatic and symptomatic individuals with familial *C9ORF72*-linked ALS-FTD (DeJesus-Hernandez et al., 2011; Renton et al., 2011).

## Results

### Selection of TDP-43-dependent cryptic exon targets

A series of human TDP-43-associated cryptic exons were identified from RNA sequencing of HeLa cells (Ling et al., 2015) and induced pluripotent stem cell (iPSC)-derived motor neurons (Klim et al., 2019; Melamed et al., 2019) depleted of TDP-43 using siRNA. Some of these cryptic exons were selected as targets (Fig. 1A) for development of monoclonal antibodies based on the following criteria: 1) the cryptic exon is spliced in-frame without premature termination codons; 2) the cryptic exon-containing gene is ubiquitously expressed or expressed abundantly in the central nervous system (CNS); and 3) the cryptic exon-encoded peptide contains immunogenic epitopes.

**Figure 1:**
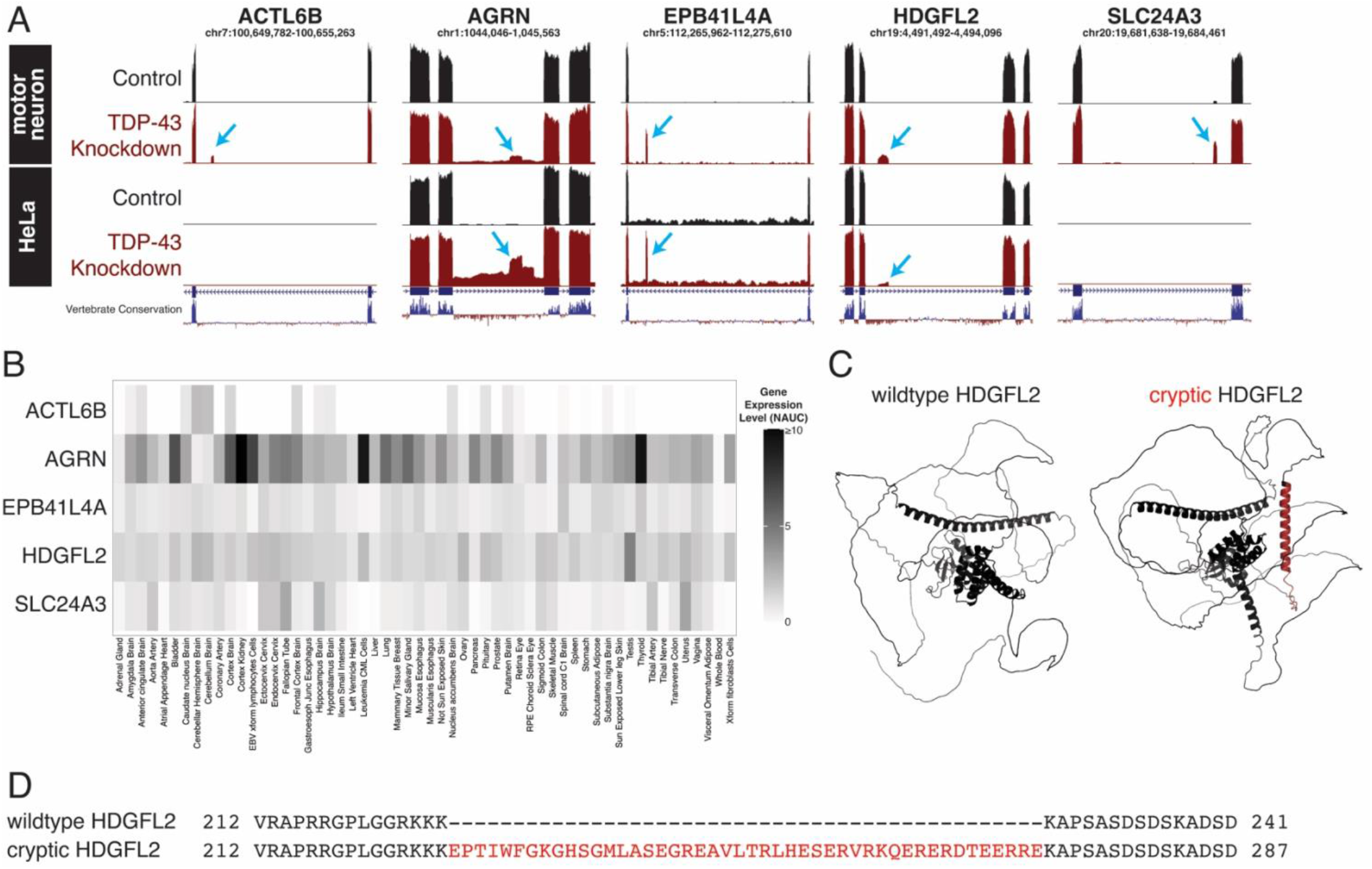
Identification of human in-frame TDP-43-associated cryptic exons. (A) UCSC Genome Browser visualization of selected cryptic exons in human motor neurons (Klim, JR *et al*. 2019) and HeLa cells (Ling, JP *et al*. 2015) aligned to the GRCh38 assembly. Red tracks indicate TDP-43 knockdown, and blue arrows identify the nonconserved cryptic exons. Gene annotation below the RNA-sequencing tracks indicate the canonical exons (thick lines) and introns (thin lines). (B) Visualization of tissue-type specific gene expression of *ACTL6B*, *AGRN*, *EPB41L4A*, *HDGFL2*, and *SLC24A3*. While *AGRN* and *HDGFL2* RNA transcripts are ubiquitously expressed, *ACTL6B*, *EPB41L4A*, and *SLC24A3* are expressed in a more tissue-specific manner. Normalized area under the curve (NAUC) values from ASCOT (Ling, JP *et al*. 2020) are used to approximate gene expression levels in different human tissue types. (C) Comparison of wild-type (left) and cryptic (right) HDGFL2 protein structures. Inclusion of the cryptic exon in mRNA leads to the addition of 46 amino acids predicted to form an alpha helix structure (red) between flanking unstructured regions. Both structures are generated using AlphaFold predictions derived from amino acid sequences. Wild-type HDGFL2 protein structure can be found on the AlphaFold protein structure database (UniProt: Q7Z4V5). (D) Alignment of wild-type and cryptic HDGFL2 amino acid sequences. Cryptic inclusion is 46 amino acids long (red) and does not impact flanking amino acids.

Genes harboring TDP-43-related cryptic exons were analyzed using alternative splicing catalog of the transcriptome (ASCOT) (Ling et al., 2020). In-frame cryptic exons in genes with high expression across different human tissues were selected for broad relevance to TDP-43-related diseases. In-frame cryptic exons in genes with high expression in the CNS were also selected due to expected involvement in ALS-FTD (Fig. 1B). Following selection of cryptic exon targets based on the initial two criteria, AlphaFold protein structure prediction software (Jumper et al., 2021) was used to model the cryptic peptide-containing proteins of interest to visualize the accessibility of the cryptic epitopes and predict the preservation of the proteins’ native conformations (Fig. 1C). Cryptic exon-encoded epitopes with immunogenic potential that did not significantly disrupt protein conformation or interfere with binding of antibodies targeting wild-type sequences were selected as prospective targets.

Of the cryptic exons meeting these criteria, one promising target selected for development of novel monoclonal antibodies was the cryptic exon-encoded epitope within Hepatoma-Derived Growth Factor-Like protein 2 (HDGFL2), a histone-binding protein that is nearly ubiquitously expressed and is detected in brain and spinal motor neurons (Fig. 1B). Mice were immunized with the *HDGFL2* cryptic exon-encoded peptide (Fig. 1D), antibody-secreting hybridoma cells were isolated for this target, and antibodies were purified from hybridoma cell lines.

### Monoclonal antibody specific to human TDP-43-dependent cryptic exon-encoded neo-epitope

A three-part screening approach was used to evaluate the sensitivity and specificity of the novel monoclonal antibodies. We first screened for monoclonal lines that would recognize the cryptic exon-encoded peptide in HDGFL2 (termed cryptic HDGFL2) when expressed as a myc-tagged green fluorescent protein (GFP)-cryptic exon fusion protein. Lysates from HEK293 cells transfected with either the GFP-myc-cryptic HDGFL2 fusion or GFP alone were subjected to protein blot analysis with monoclonal antibodies. Of monoclonal lines #1-65 through 1-71 against the cryptic HDGFL2 epitope, lines #1-66 and 1-69 detected with specificity the fusion protein containing the cryptic HDGFL2 peptide (Fig. S2). Line #1-69 was used moving forward.

Second, we used an siRNA knockdown strategy in HeLa cells to deplete TDP-43 (siTDP HeLa) and screened antibodies in siTDP vs. control HeLa lysates (Fig. 2A). We confirmed, as expected, the appearance of *HDGFL2* containing cryptic exons in siTDP compared to control cells (Fig. 2B). We then subjected total cell extracts from control and siTDP HeLa cells to protein blot analysis with the monoclonal antibodies to determine specificity of the antibodies. As expected, a band corresponding to the normal HDGFL2 protein was observed in both control and siTDP HeLa cells using an antibody recognizing the native HDGFL2 (Fig. 2C). Importantly, the monoclonal #1-69 cryptic antibody recognized a novel band of the expected size, presumably corresponding to the cryptic peptide-containing protein, in extract of HeLa cells depleted of TDP-43 but not control cells (Fig. 2C). To further demonstrate the specificity of the monoclonal antibody, we performed immunoprecipitation (IP) using this antibody followed by protein blot analysis with the antibody recognizing the native HDGFL2. While no positive band of correct molecular weight was observed in control extract, a robust band of expected size for the cryptic exon-encoded peptide within HDGFL2 was evident in siTDP HeLa cells (Fig. 2D). Thus, these data demonstrate that this monoclonal antibody specifically recognizes the cryptic exon-encoded peptide within HDGFL2.

**Figure 2.**
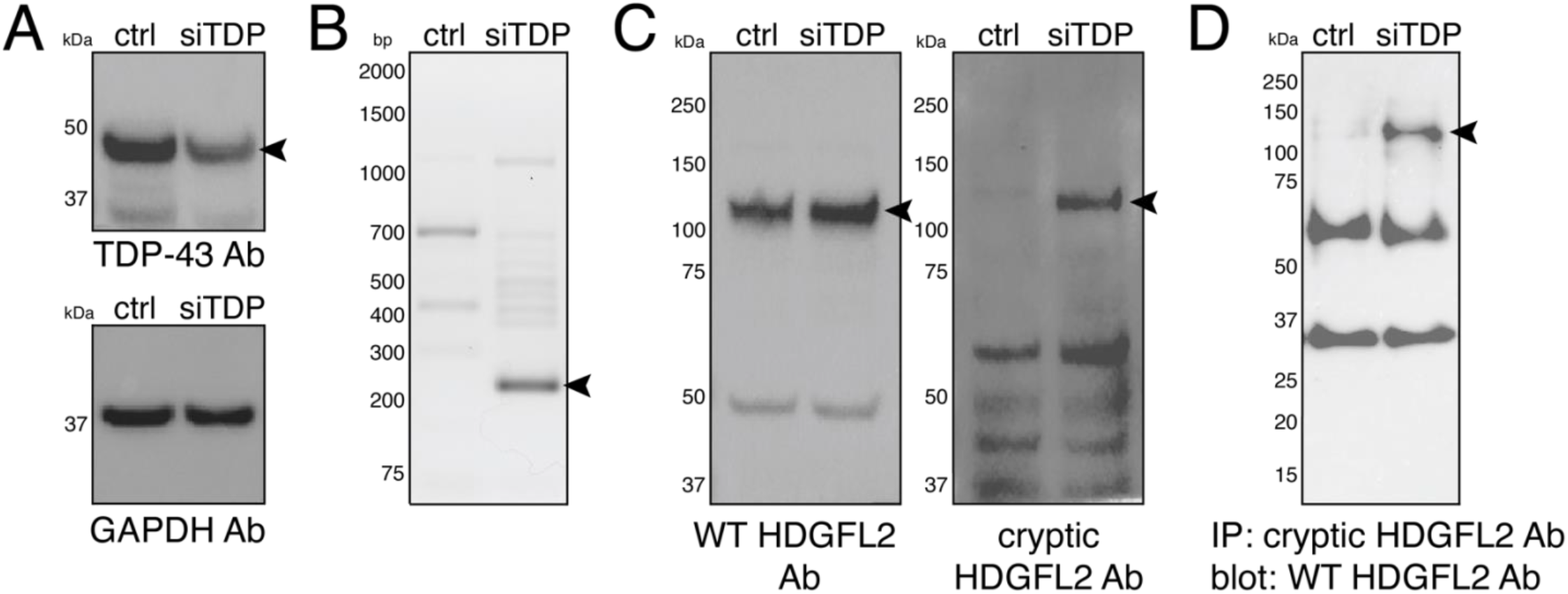
Novel antibody shows specificity for HDGFL2 with cryptic peptide. (A) TDP-43 is reduced in HeLa treated with TDP-43 siRNA (siTDP). (B) RT-PCR with primers designed to amplify the cryptic exon sequence of HDGFL2 shows a product only in siTDP. (C) Protein extracts as in panel A were subjected to protein blot analysis using an antibody against the native HDGFL2 protein (left) or the novel monoclonal antibody (1-69) against the cryptic sequence in HDGFL2 (right). HDGFL2 harboring the neo-epitope is only detected in siTDP. (D) IP blot using 1-69 cryptic antibody for pulldown and WT HDGFL2 antibody for blotting reveals a band of the expected size only in siTDP.

### A highly sensitive MSD sandwich ELISA for detection of TDP-43-dependent cryptic peptide

To employ our novel monoclonal antibodies to detect cryptic exon-encoded peptides in human CSF, we developed a highly sensitive sandwich ELISA using the Meso Scale Discovery (MSD) platform and validated this assay for the cryptic exon target in HDGFL2. In the same manner as we performed IP-protein blot analysis, we used the #1-69 cryptic monoclonal antibody as the capture (coating) antibody to pull down the cryptic exon-encoded peptide within HDGFL2 and the rabbit antibody recognizing the wild-type (WT) HDGFL2 as the primary detection antibody. The amount of cryptic HDGFL2 can be quantified indirectly by using a sulfo-tagged anti-rabbit antibody for secondary detection (Fig. 3A).

**Figure 3.**
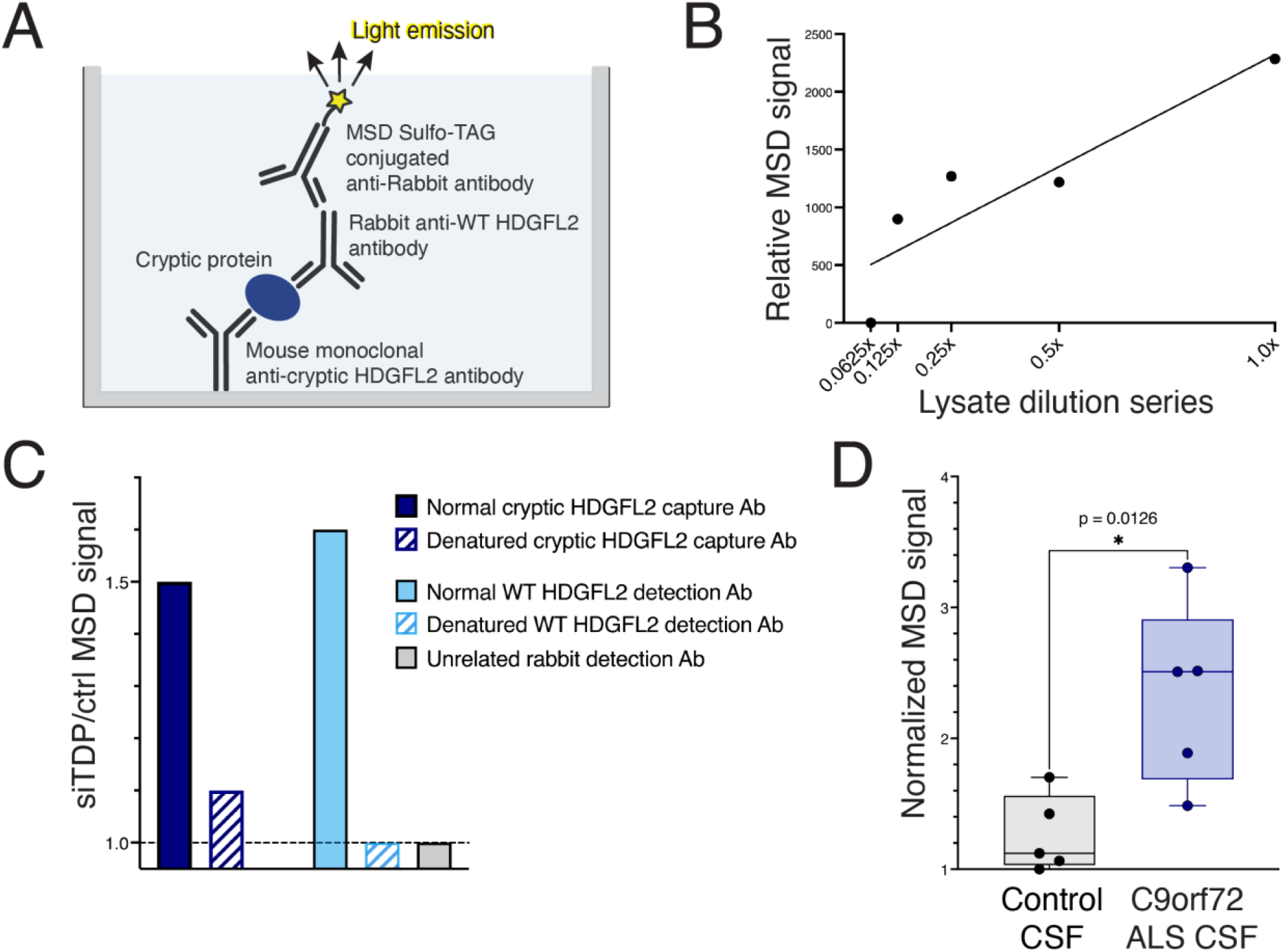
Development of an MSD assay specific for cryptic HDGFL2. (A) Sandwich ELISA using Meso Scale Discovery (MSD) system. (B) Dose-dependent increase of MSD signal in siTDP HeLa lysate compared to control HeLa lysate. MSD signal of control HeLa is subtracted from that of siTDP. (C) Elevated siTDP MSD signal is specific to intact capture and detection antibodies. (D) Average cryptic HDGFL2 signal is elevated in CSF of *C9ORF72* ALS compared to controls.

We tested this MSD ELISA protocol using the control and siTDP HeLa lysates. Compared to control lysate, a concentration-dependent increase in MSD signal was observed in siTDP HeLa lysate (Fig. 3B). When the primary detection antibody against WT HDGFL2 was replaced with an unrelated rabbit antibody, no difference between MSD signal of siTDP and control HeLa lysates was observed (Fig. 3C), indicating that the siTDP signal is specific. When the primary detection antibody against WT HDGFL2 was denatured by heating at 95°C for 30 minutes prior to addition to the plate, there was again no difference between MSD signal of siTDP and control HeLa lysates (Fig. 3C). These data demonstrate the specificity of the assay for HDGFL2 binding by the detection antibody. When the cryptic HDGFL2 capture antibody was denatured by heating at 95°C for 30 minutes prior to addition to the plate, the ratio of siTDP/control HeLa MSD signal decreased to control level (Fig. 3C). These data support the specificity of the assay for cryptic HDGFL2 binding by the capture antibody. Taken together, these data establish a highly sensitive and specific sandwich ELISA for the detection of cryptic peptide-containing protein, a valuable method to monitor TDP-43 loss of function in biofluids of ALS patients.

### Presence of cryptic HDGFL2 in CSF of pre-symptomatic *C9ORF72* mutation carriers

In a pilot study using CSF of *C9ORF72* mutation carriers with ALS and control individuals, the MSD signal for cryptic HDGFL2 was significantly elevated in ALS samples (mean=1601) as compared to those of controls (mean=−64.10; p<0.013; Fig. 3D). We then expanded our study into a larger, longitudinal cohort of 46 *C9ORF72* mutation carriers with a total of 88 CSF samples, with each individual contributing one, two, or three time points of CSF (Traynor et al., 2000). This cohort of *C9ORF72* mutation carriers offered a unique opportunity to assay pre-symptomatic individuals, as subjects were recruited due to their mutation status, but not all had phenoconverted to symptomatic disease yet.

The study cohort included 31 individuals with symptomatic ALS (n=20), ALS-FTD (n=8), or FTD (n=3) and 15 pre-symptomatic individuals. The mean age was 52.0 years old (range=28.6-72.6, standard deviation [SD]=11.4), and 47.8% (22/46) were female.

Normalized cryptic HDGFL2 MSD signals in males (mean=759.1) were significantly higher compared to females (mean=689.0, two sample t-test, p=0.03). However, there was no difference between cryptic HDGFL2 MSD signals in symptomatic males (mean=769.9) and symptomatic females (mean=741.4, two sample t-test, p=0.52).

Normalized cryptic HDGFL2 signals in both pre-symptomatic (mean=665.4) and symptomatic (mean=758.4) *C9ORF72* mutation carriers were significantly higher compared to controls (mean=451.4) (one-way ANOVA with Tukey’s multiple comparisons test, Fig. 4A). We also assayed CSF of 6 sporadic ALS cases. While cryptic HDGFL2 signals (mean=513.0) were elevated compared to controls, this difference was not significant.

**Figure 4.**
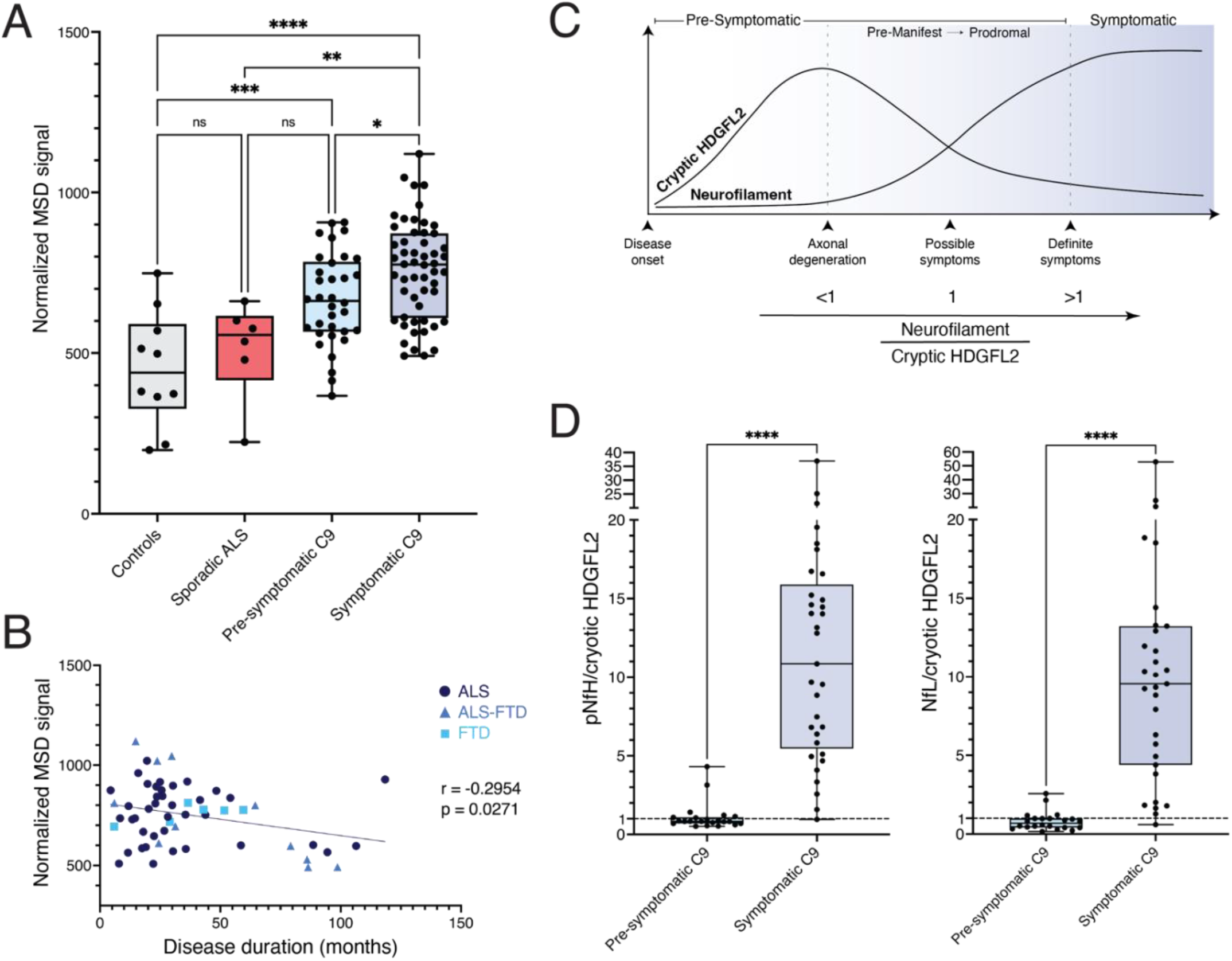
Cryptic HDGFL2 can be detected by MSD assay in CSF of both pre-symptomatic and symptomatic *C9ORF72* mutation carriers. (A) Elevated cryptic HDGFL2 is detected in pre-symptomatic and symptomatic *C9ORF72* mutation carriers compared to controls. (B) Cryptic HDGFL2 detection by our MSD assay in CSF of *C9ORF72* mutation carriers diagnosed with ALS, FTD, or ALS-FTD tends to be higher during the earlier stage of symptomatic disease. Pearson correlation, *r* = −0.30, *p* = 0.027. (C) ALS staging model (adapted from Benatar et al., 2019) based on the dynamic of neurofilament subunit and cryptic HDGFL2 accumulation in CSF of ALS patients. As our preliminary data indicate that a trend towards higher levels of cryptic HDGFL2 is observed during earlier-stage disease, we hypothesize that while CSF neurofilament subunits rise during the prodromal phase and remain elevated in symptomatic disease, CSF cryptic HDGFL2 would decrease after its peak, which may occur during the prodromal stage. Thus, a ratio of pNfH or NfL to cryptic HDGFL2 of less than 1 or greater than 1 would indicate, respectively, pre-symptomatic or symptomatic phase of disease. (D) Ratios of pNfH/cryptic HDGFL2 and NfL/cryptic HDGFL2 in CSF of presymptomatic and symptomatic *C9ORF72* mutation carriers. Using the ratio of pNfH and NfL to cryptic HDGFL2 in CSF, it may be possible to establish an ALS staging scale spanning from pre-symptomatic phase (ratio <1) to symptomatic phase (ratio >1).

*C9ORF72* mutation carriers&#8217; levels of cryptic HDGFL2 did not correlate with their scores on the Revised Amyotrophic Lateral Sclerosis Functional Rating Scale (ALSFRS-R) (Pearson correlation, r=− 0.11, p=0.31) (Cedarbaum et al., 1999). Additionally, cryptic HDGFL2 signals did not correlate with rate of disease progression as measured by change in ALS-FRS-R score per months elapsed since the prior CSF collection point (Fig. S3; Pearson correlation, r=−0.12, p=0.52). However, among the 31 individuals who had phenoconverted to symptomatic ALS, ALS-FTD, or FTD, levels of cryptic HDGFL2 showed a negative correlation with symptom duration (Pearson correlation, r=−0.30, p=0.03; Fig. 4B), suggesting that levels of cryptic HDGFL2 tend to be higher during the earlier stage of symptomatic disease.

### CSF ratio of pNfH or NfL/cryptic HDGFL2 is a marker for phenoconversion

Cryptic HDGFL2 appears to increase during the pre-symptomatic stage of disease and may decrease with symptomatic disease progression, and this trend is in the direction opposite to that of phosphorylated neurofilament heavy (pNfH) and neurofilament light (NfL) chains (Benatar et al., 2018, 2019). Therefore, we analyzed the ratios of pNfH and NfL to cryptic HDGFL2 (pNfH/cryptic HDGFL2, NfL/cryptic HDGFL2), anticipating that a ratio less than 1 may indicate a pre-symptomatic stage of disease, while a ratio greater than 1 would indicate a symptomatic stage (Fig. 4C). For each of the longitudinal samples, we determined the ratios of pNfH and NfL to cryptic HDGFL2. For the pre-symptomatic group, 77% (17/22) of pNfH/cryptic HDGFL2 ratios were less than or equal to 1, and 82% (18/22) of NfL/cryptic HDGFL2 ratios were less than 1. Of note, the two highest ratios of both pNfH/cryptic HDGFL2 and NfL/cryptic HDGFL2 for the pre-symptomatic group were identified in one individual whose ALS-FRS-R score (Cedarbaum et al., 1999) had fallen from 47 to 43 despite still being categorized as “asymptomatic.” For the symptomatic group, 97% (32/33) of pNfH/cryptic HDGFL2 ratios and 97% (30/31) of NfL/cryptic HDGFL2 ratios were greater than 1 (Fig. 4D). These data support the idea that the ratio of neurofilament subunits/cryptic HDGFL2 in CSF of *C9ORF72* ALS patients is a predictor of phenoconversion.

## Discussion

Previously, we proposed that loss of TDP-43 splicing repression of cryptic exons underlies disease pathogenesis of ALS-FTD (Ling et al., 2015). This view is further supported by identification of key cryptic exon targets of TDP-43 (Brown et al., 2022; Klim et al., 2019; Ma et al., 2022; Melamed et al., 2019; Prudencio et al., 2020). However, evidence that such loss of TDP-43 function occurs during early-stage disease, rather than being an end-stage phenomenon, remains elusive, as no method to detect TDP-43 dysfunction in living individuals currently exists.

Our findings demonstrating the presence of cryptic HDGFL2 in CSF of ALS patients now provide direct evidence that loss of TDP-43 splicing repression occurs during early-stage disease, including the pre-symptomatic phase (Fig. 4). These findings align with a case in which a patient underwent surgical resection of her temporal lobe for treatment of epilepsy 5 years before developing FTD symptoms. Her resected brain tissue was shown to have lost nuclear TDP-43 in some neurons but not contain any cytoplasmic inclusions, supporting the notion that loss of TDP-43 splicing repression represents an early event that drives neuron loss (Vatsavayai et al., 2016). Our detection of TDP-43-dependent cryptic peptides in CSF of pre-symptomatic *C9ORF72* mutation carriers also aligns with a common theme of neurodegenerative diseases&—that pathogenic mechanisms often begin years before clinically relevant symptoms emerge.

Our analysis of cryptic exon accumulation in CSF of numerous *C9ORF72*-linked individuals allows us not only to establish loss of TDP-43 splicing repression occurring at a pre-symptomatic stage, but also to discern the dynamic nature of TDP-43 cryptic proteins in which the ratio of neurofilament heavy or light chains to cryptic HDGFL2 can be employed as a biomarker that may signal symptom onset of ALS. We thus propose a biomarker trajectory whereby the ratio of pNfH or NfL/cryptic HDGFL2 is less than 1 during a pre-symptomatic stage of disease and increases to greater than 1 as symptom onset occurs, suggesting the ability to predict phenoconversion in familial ALS.

While further study of individuals with sporadic ALS is warranted, these findings suggest that cryptic exon-encoded peptides could serve as biomarkers to facilitate earlier diagnosis of ALS, which is currently delayed by the need to rule out mimic conditions (Traynor et al., 2000). Such early-stage biomarkers that can identify ALS-related pathomechanisms are currently critically lacking. Earlier diagnosis would permit prompt therapeutic intervention with a greater chance of success.

Additionally, our findings that loss of TDP-43 splicing repression occurs during the pre-symptomatic stage of disease provide strong rationale to develop therapeutic strategies to complement TDP-43 splicing repression deficits for ALS—for example, an AAV9 gene therapeutic strategy that expresses a splicing repressor, termed CTR (Donde et al., 2019). Because detection of cryptic exon-encoded peptides in patient biofluids reflects loss of TDP-43 function, evaluating the dynamics of cryptic peptide biomarkers could provide a way of measuring target engagement for new therapeutics aimed at restoring TDP-43 function. Many clinical trials for ALS treatments currently suffer from an inability to measure therapeutic efficacy at a mechanistic level (Mitsumoto et al., 2014), so cryptic peptide biomarkers could be used in the future to better understand the results of clinical trials and improve drug design.

While there is some evidence that NfL and pNfH levels may rise pre-symptomatically in ALS (Benatar et al., 2018, 2019), these markers reflect broad neurodegeneration and are elevated in many diseases, limiting their diagnostic utility, and they do not reflect TDP-43 dysfunction, limiting their ability to measure target engagement in clinical trials. Additionally, *C9ORF72*-related biomarkers have been studied, such as dipeptide repeats (Balendra et al., 2017; Krishnan et al., 2022), but the functional relevance of these proteins is still unclear. Thus, cryptic peptide biomarkers would transform our current biomarkers of ALS-FTD through their specificity for TDP-43-related disease and their reflection of TDP-43 dysfunction.

Furthermore, involvement of TDP-43 dysfunction early in *C9ORF72* ALS suggests that the current standard of detecting TDP-43 pathology in postmortem tissues may miss cases of TDP-43 misregulation. TDP-43 pathology is identified by cytoplasmic inclusions reactive to anti-TDP-43 or anti-phosphorylated TDP-43 antibodies. However, a lack of overt cytoplasmic staining of TDP-43 aggregates does not rule out loss of TDP-43 splicing repression. Inclusion of cryptic exons early in disease indicates that identifying TDP-43 nuclear depletion or TDP-43-related cryptic proteins in histological sections may provide a more sensitive approach to classification of TDP-43 pathology.

In the future, we plan to develop a highly sensitive multiplexed MSD assay so that we can measure cryptic HDGFL2 and other disease-relevant markers, such as pNfH, NfL, tau, amyloid-β, and α-synuclein simultaneously, providing additional insight into disease staging for ALS and other neurodegenerative diseases exhibiting TDP-43 pathology. This multiplexed system will also allow us to analyze several cryptic peptides at once. We envision the ability to detect a set of TDP-43 cryptic exon targets which may display different dynamics throughout disease progression. Analyzing the changes of these targets throughout disease could provide detailed information on disease staging, progression, and potentially prognosis that is not currently available for ALS-FTD.

The impact of this work also extends beyond the ALS-FTD spectrum. As many cases of Alzheimer’s disease (AD) also possess TDP-43 pathology (Josephs, Murray, et al., 2014; Josephs, Whitwell, et al., 2014; Meneses et al., 2021; Nelson et al., 2022; Robinson et al., 2018), cryptic peptide biomarkers would be helpful in distinguishing “pure AD” from mixed (or multiple) etiology dementia (MED), which likely warrant different treatment strategies. Cryptic peptide biomarkers could be used in the future to stratify MED and pure AD for clinical trials, giving therapeutics the best chance of success. They could also be used retrospectively to assess whether subgroups of prior clinical trials display different therapeutic efficacy than when analyzed as a single disease group. Several other neurological conditions involve TDP-43 proteinopathy, including LATE (Nelson et al., 2019), multiple sclerosis (Masaki et al., 2020), and chronic traumatic encephalopathy (McKee et al., 2010); therefore, the impact of these biomarkers could be far-reaching.

## Methods

### Analysis of RNA-seq data

Fastq files were downloaded from the NCBI’s Sequence Read Archive and aligned to the GRCh38 human genome assembly using STAR (version 2.7.10a) with default parameters. Megadepth was used to convert the output BAM files to BigWig files, and the data were then displayed on the UCSC Genome Browser (http://genome.ucsc.edu/) to visualize the cryptic exons.

The data table containing NAUC information was downloaded from ASCOT (Ling et al., 2020). Data was subset to the genes of interest, and a heatmap was generated using ggplot2 package in R.

### Protein structure generation and comparison

The cryptic peptide sequences were identified by translating the cryptic exon sequences. The predicted wild-type protein structures were generated using the AlphaFold Monomer v2.0 pipeline and downloaded from the AlphaFold Protein Structure Database. The predicted cryptic protein structures were generated by entering the amino acid sequences of interest into the AlphaFold Monomer v2.0 pipeline (version 2.2.0).

### Generation of monoclonal antibodies

Monoclonal antibodies were generated by CDI Laboratories, Inc. Mice were immunized with the cryptic peptide of interest, and hybridomas were produced from these mice. IgG-positive hybridomas were identified by ELISA. Hybridoma lines that produced antibodies recognizing their cognate antigen as the top target on HuProt human protein microarray, which contains >19,500 affinity-purified recombinant human proteins (Venkataraman et al., 2018), were identified. Promising hybridoma lines were further screened by protein blot.

### Generation of wild-type and cryptic *HDGFL2* expression vectors

The wild-type *HDGFL2* mRNA sequence (ENST00000616600.5) was identified using UCSC Genome Browser. RNA-sequencing visualization of TDP-43 knockdown motor neurons (Klim et al., 2019) on UCSC Genome Browser was used to extract the cryptic exon sequence. Codons corresponding to a glycine serine linker and 6-HisTag (GGGSHHHHHH) were added to the 3’ end of the sequence directly before the stop codon. A Kozak sequence was added to the 5’ end of the sequence. Codons were optimized by using IDT Codon Optimizer and modifying codons that led to nucleic acid repeats of four or more while retaining the amino acid translation. Restriction sites corresponding to NsiI, XmnI, BclI, BstXI, NheI, BspEI, XhoI, XbaI, PspOMI, BglII, NotI, BamHI were removed from the sequence. The resulting sequence was synthesized into the pTwist CMV Puro expression vector by Twist Bioscience.

### RT-PCR analysis

RNA was extracted from HeLa samples using TRIzol (Life Tech., 15596-026) and RNeasy Mini Kits (Qiagen, 74104). cDNA was derived from total RNA using ProtoScript II First Strand cDNA Synthesis Kit (NEB, E6560S). Numerous primers were designed against cryptic exon targets and screened to identify primer pairs that minimized background bands.

### Protein blot and Immunoprecipitation-protein blot analysis

Immunoprecipitation was performed by overnight incubation of HeLa lysates with the novel #1-69 antibody against cryptic HDGFL2 at 4°C with rotation. Then a 50% slurry of Protein G agarose beads (Cell Signaling, #37478) in RIPA lysis buffer was added, and this mixture was incubated with rotation at 4°C for 1-3 hours. Bead complexes were denatured in NuPAGE LDS Sample Buffer (4X) at 70°C for 10 minutes and microcentrifuged at 14,000 g for 1 minute. Samples were then analyzed by protein blot.

Protein blot analysis was performed following electrophoresis using NuPAGE 4-12% Bis-Tris polyacrylamide gels. Proteins were transferred onto PVDF membranes using the iBlot™ Dry Blotting System from Invitrogen. After blocking of the membrane, primary antibodies were incubated overnight at 4°C with rocking.

### Meso Scale Discovery ELISA

For our novel sandwich ELISA using the Meso Scale Discovery (MSD) platform, our novel antibody (#1-69) against the cryptic exon-encoded peptide target in HDGFL2 served as the capture antibody, and a commercial antibody against the wild-type protein was used as the primary detection antibody (Anti-CTB-50L17.10 antibody produced in rabbit; Prestige Antibodies® Powered by Atlas Antibodies). A species-specific sulfo-tagged antibody (Anti Rabbit Antibody Goat SULFO-TAG Labeled; Meso Scale Discovery) was used as a secondary detection reagent to generate electrochemiluminescence. Assays were conducted on MSD MULTI-ARRAY 96-well SECTOR plates and measured with the MESO QuickPlex SQ 120 MM instrument.

An original pilot study of five *C9ORF72*-associated ALS and five control subjects utilized duplicates of 120 microliters (μL) of CSF for each individual. In the following studies, CSF samples were assayed with 25 μL of CSF diluted in 75 μL of MSD diluent 35, for a total of 100 μL added per well. A standard curve was constructed using a concentration series of lysate from HeLa cells transfected with the cryptic *HDGFL2* expression vector (Fig. S4). This curve was employed on each plate to normalize signals based on a four-parameter logistic (4PL) fit. In longitudinal assays, all time points of CSF collection for one individual were assayed on the same plate. Immediately before use, CSF was thawed, vortexed, and briefly spun down. Unless otherwise stated, *C9ORF72* CSF used was freshly thawed without any previous freeze-thaw cycles. Upon normalization, one *C9ORF72* CSF sample was below the lower limit of detection and thus was excluded from further analyses. Any samples previously thawed one time were normalized based on the cryptic HDGFL2 signal differences between a previously thawed aliquot and a previously unthawed aliquot of a set of *C9ORF72* CSF samples (Fig. S5).

### Cerebrospinal Fluid Samples

CSF samples from *C9ORF72* mutation carriers were provided by the Natural History and Biomarker Study of *C9ORF72* ALS (protocol 13-N-0188) with enrollment/study period: 2013-2020 (Offit et al., 2020). CSF samples from sporadic ALS subjects and from healthy and non-ALS neurological disease control subjects were provided by the Northeast Amyotrophic Lateral Sclerosis (NEALS) Biorepository and Biogen, Inc./Precision Med. Additional control CSF samples were obtained from individuals with hydrocephalus at Johns Hopkins Bayview Medical Center. Phosphorylated neurofilament heavy chain (pNfH) levels in CSF were measured by Biogen using the ProteinSimple Ella microfluidic immunoassay according to the manufacturer’s instructions. Neurofilament light chain (NfL) levels in CSF were measured by Biogen using the Quanterix Simoa assay.

### Statistical Analysis

Statistics were performed in Stata 17 and GraphPad Prism.

### Key Resources Table

| **Antibody target** | **Source** | **Identifier** |
| --- | --- | --- |
| **Western Blot** |  |  |
| Wild-type HDGFL2 | Atlas Antibodies | HPA044208 |
| TDP-43 | Proteintech | 10782-2-AP |
| GAPDH | Cell Signaling Technology | 5174 |
| **MSD ELISA** |  |  |
| Wild-type HDGFL2 | Atlas Antibodies | HPA044208 |
| Rabbit IgG (sulfo-tagged) | Meso Scale Discovery | Anti Rabbit Antibody Goat<br>SULFO-TAG Labeled |

### Data availability

RNA-seq data analyzed is available on NCBI’s Sequence Read Archive under SRA study numbers SRP166282 (Klim et al., 2019) and SRP057819 (Ling et al., 2015). NAUC data tables are available on http://ascot.cs.jhu.edu/ (Ling et al., 2020).

## Acknowledgments

We acknowledge the NEALS Biorepository for providing all or part of the biofluids from the ALS, healthy controls, and non-ALS neurological controls used in this study. This work was supported in part by the NINDS, NIH (R01NS095969 to PCW), the Robert Packard Center for ALS Research at Johns Hopkins (to PCW), the Target ALS Foundation (to PCW), and the Karen Toffler Charitable Trust (to KI).

Part of this work was carried out at the Advanced Research Computing at Hopkins (ARCH) core facility (rockfish.jhu.edu), which is supported by the National Science Foundation (NSF) grant number OAC 1920103.

## Author contributions

KEI, JPL, and PCW conceptualized study; KEI and PCW wrote manuscript, and all other authors edited manuscript; KEI, PJ, KEB, DR, DB, IS, and KDB performed experiments and analyzed data; BT, JDB, DR, DB, AM, and ESO provided CSF samples.

---

**Figure S1.**
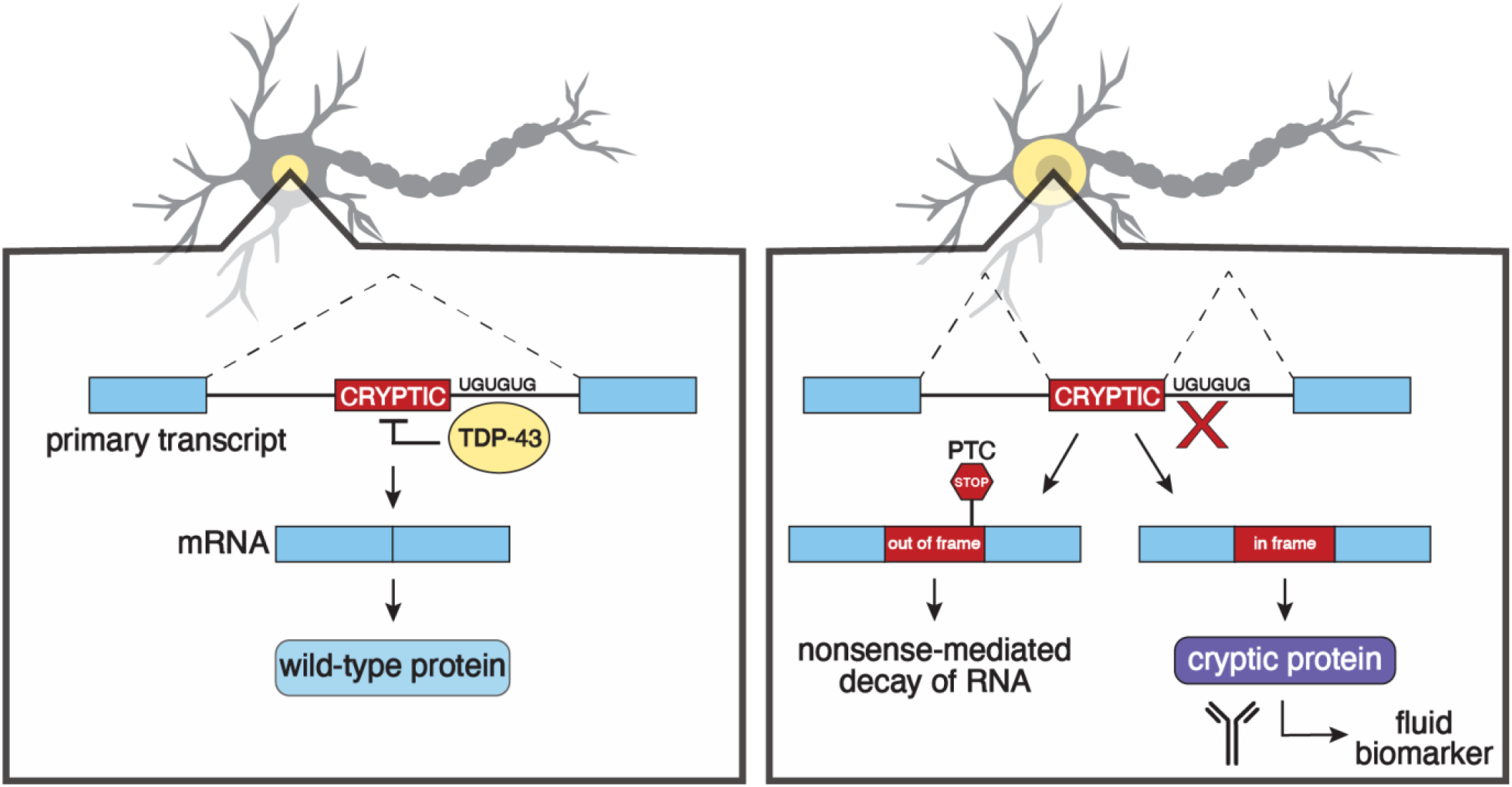
Strategy for developing cryptic peptide fluid biomarkers. When TDP-43 is lost from the nucleus, it fails to repress the splicing of cryptic exons. As some cryptic exons are incorporated in-frame, antibodies can be developed against cryptic exon-encoded peptides to serve as fluid biomarkers. (PTC=premature termination codon).

**Figure S2.**
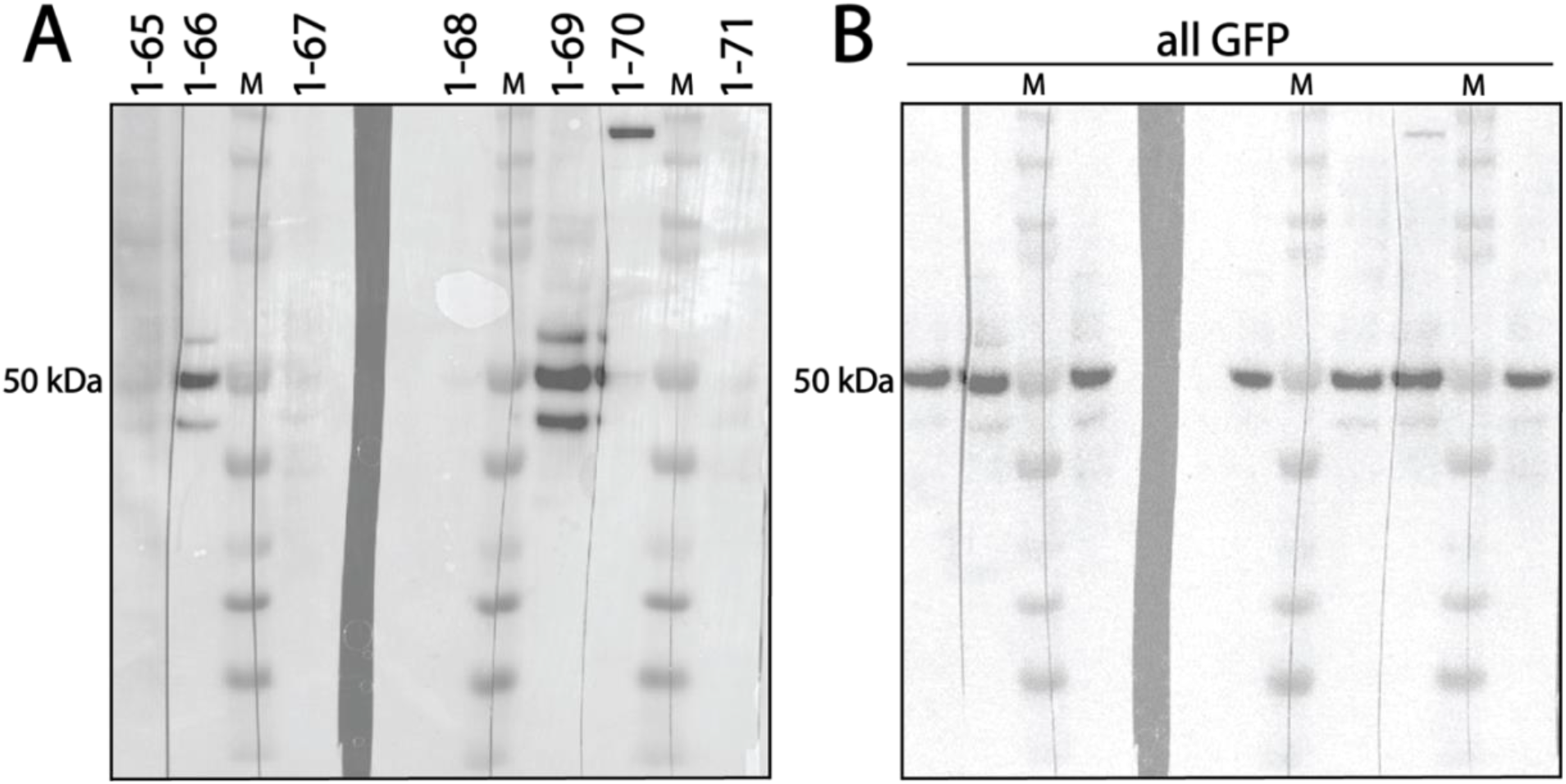
Generation of a series of antisera recognizing a cryptic peptide in HDGFL2. (A) HEK293 cells were transfected with GFP-myc-cryptic HDGFL2, and lysates were subjected to protein blot analysis using antisera from 7 different monoclonal lines against cryptic HDGFL2. Lines 1-66 and 1-69 detected the fusion protein containing the cryptic HDGFL2 peptide. (B) Immunoblots of the same lysates using anti-GFP antisera confirmed the identity of the GFP-myc-cryptic HDGFL2 fusion protein. M = molecular weight.

**Figure S3.**
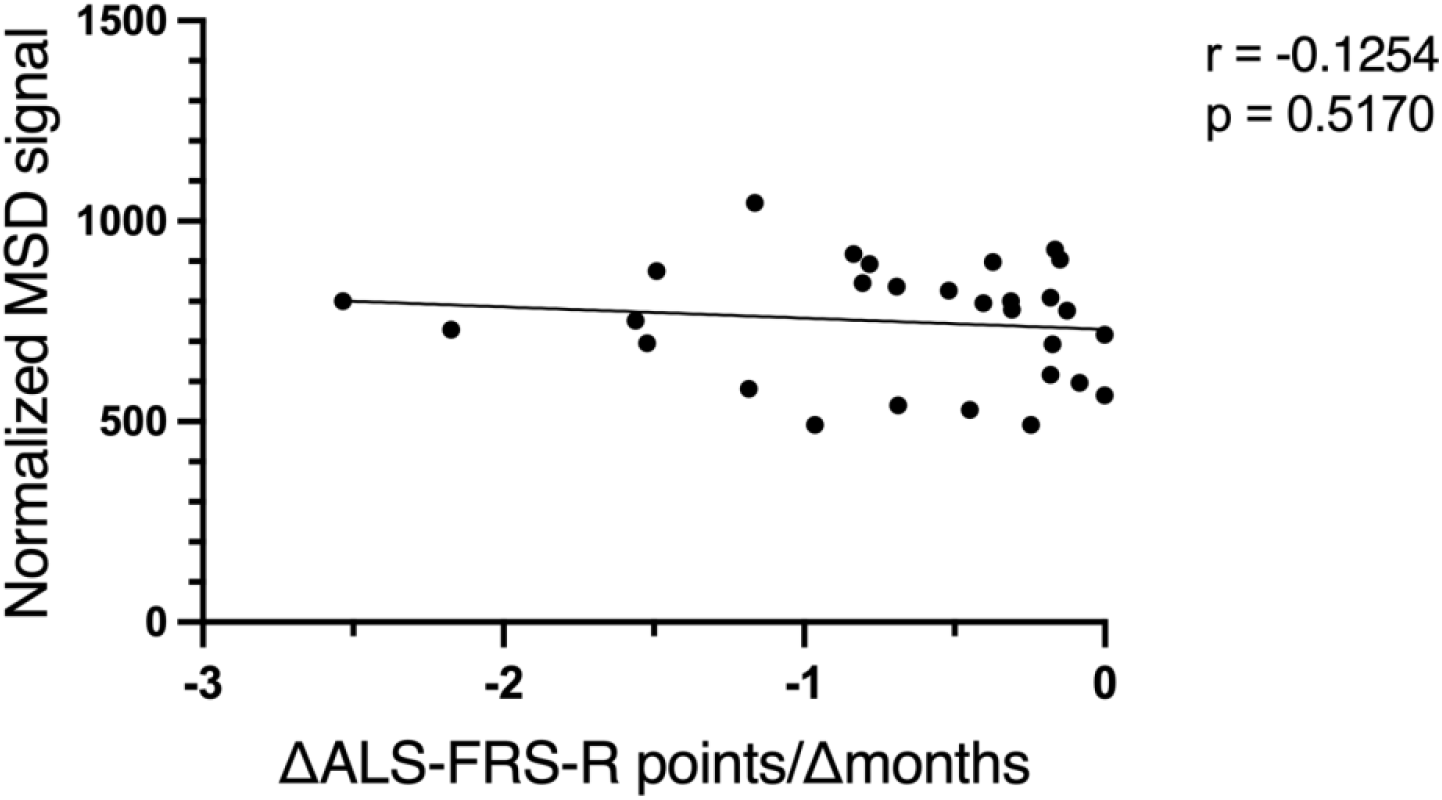
Rate of disease progression does not correlate with CSF cryptic HDGFL2 measured by MSD assay in *C9ORF72* mutation carriers. Rate of disease progression was measured as the change in ALS-FRS-R points divided by the number of months that had passed since the prior CSF collection point.

**Figure S4.**
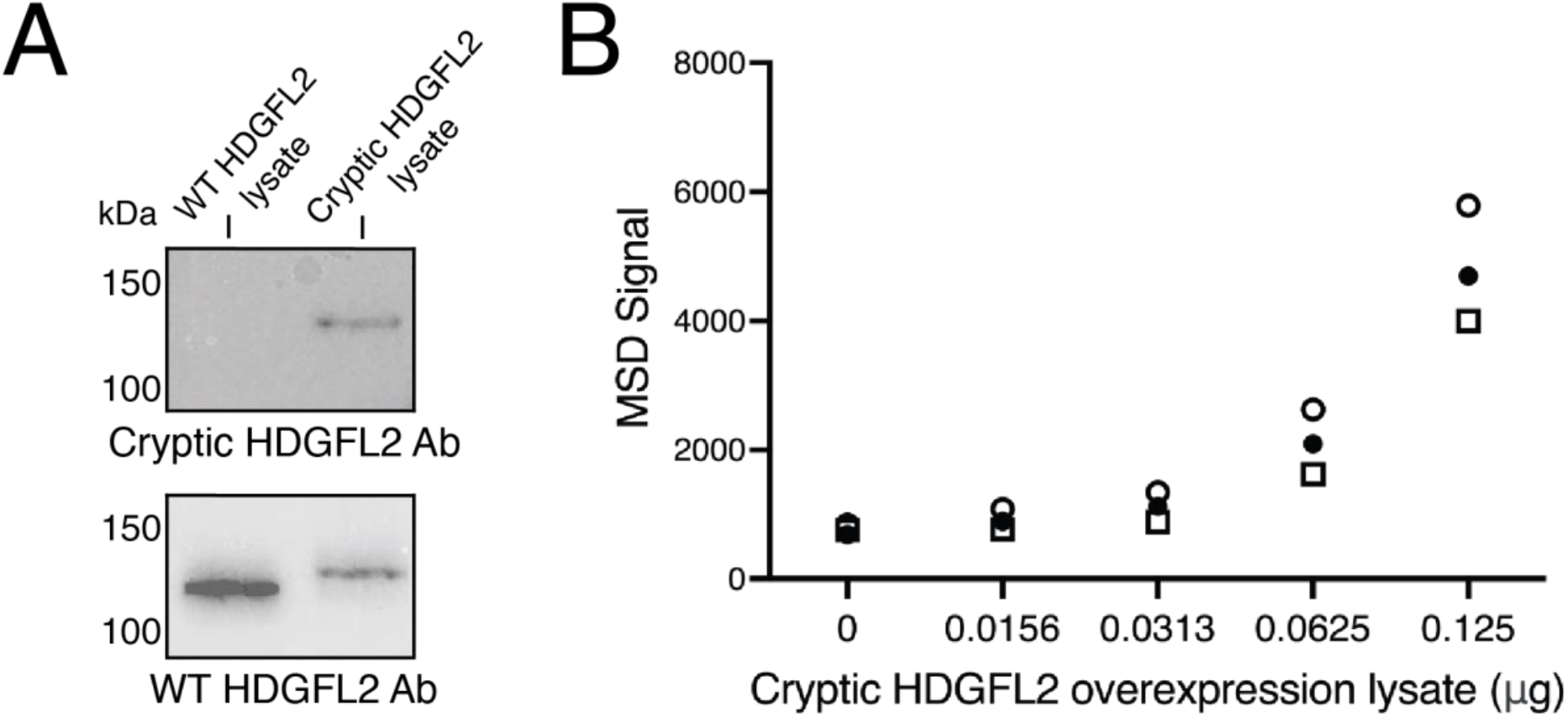
Standard curves used for normalization of signal between MSD plates. (A) HeLa cells were transfected with a construct expressing either wild-type HDGFL2 or cryptic HDGFL2. Lysate of HeLa cells overexpressing WT HDGFL2 and lysate of HeLa cells overexpressing cryptic HDGFL2 show a band of the expected size when probed with the WT HDGFL2 antibody. Only lysate of HeLa cells overexpressing cryptic HDGFL2 show a band of the expected size when probed with the novel cryptic HDGFL2 antibody. (B) Lysates of HeLa cells transfected to overexpress the cryptic HDGFL2 protein were measured in a concentration series on the three MSD plates used for the CSF assays in this study, and these standard curves were analyzed by a four-parameter logistic (4PL) fit to normalize signal between plates.

**Figure S5.**
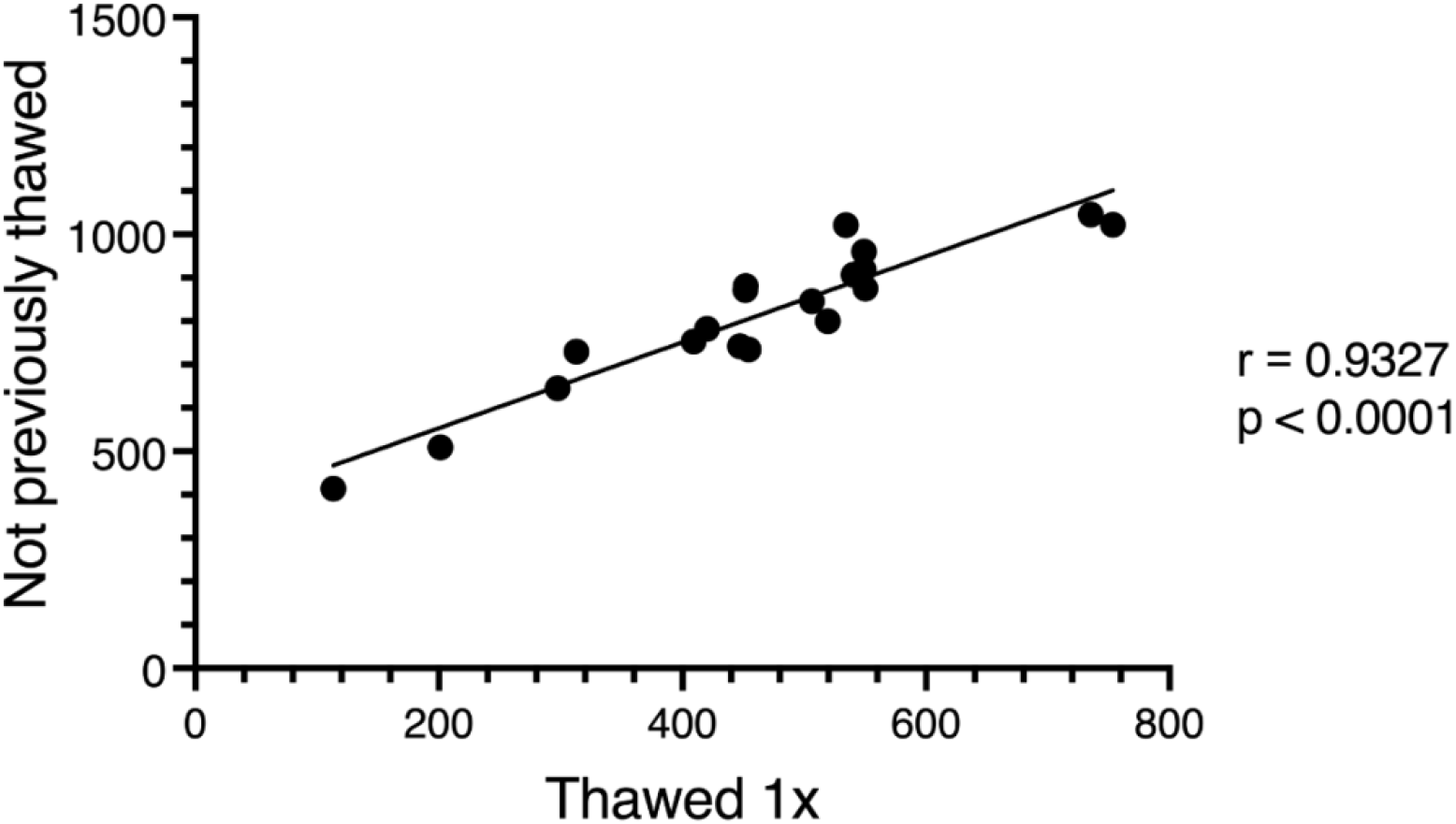
Reproducibility of cryptic HDGFL2 MSD assay. Nineteen *C9ORF72* mutation carrier CSF samples were assayed on two different days. One set of samples had not been previously thawed, while the other set had been thawed and re-frozen one time. Normalized MSD signals were slightly decreased for the previously thawed samples, but relative signal levels were preserved (r=0.93; p<0.0001).

## References

Balendra, R., Moens, T. G., & Isaacs, A. M. (2017). Specific biomarkers for C9orf72 FTD/ALS could expedite the journey towards effective therapies. EMBO Molecular Medicine, 9(7), 853–855. https://doi.org/10.15252/emmm.201707848

Barmada, S. J., Skibinski, G., Korb, E., Rao, E. J., Wu, J. Y., & Finkbeiner, S. (2010). Cytoplasmic Mislocalization of TDP-43 Is Toxic to Neurons and Enhanced by a Mutation Associated with Familial Amyotrophic Lateral Sclerosis. Journal of Neuroscience, 30(2), 639–649. https://doi.org/10.1523/JNEUROSCI.4988-09.2010

Benatar, M., Wuu, J., Andersen, P. M., Lombardi, V., & Malaspina, A. (2018). Neurofilament light: A candidate biomarker of presymptomatic amyotrophic lateral sclerosis and phenoconversion. Annals of Neurology, 84(1), 130–139. https://doi.org/10.1002/ana.25276

Benatar, M., Wuu, J., Lombardi, V., Jeromin, A., Bowser, R., Andersen, P. M., & Malaspina, A. (2019). Neurofilaments in pre-symptomatic ALS and the impact of genotype. Amyotrophic Lateral Sclerosis & Frontotemporal Degeneration, 20(7–8), 538–548. https://doi.org/10.1080/21678421.2019.1646769

Brettschneider, J., Del Tredici, K., Toledo, J. B., Robinson, J. L., Irwin, D. J., Grossman, M., Suh, E., Van Deerlin, V. M., Wood, E. M., Baek, Y., Kwong, L., Lee, E. B., Elman, L., McCluskey, L., Fang, L., Feldengut, S., Ludolph, A. C., Lee, V. M.-Y., Braak, H., & Trojanowski, J. Q. (2013). Stages of pTDP-43 pathology in amyotrophic lateral sclerosis. Annals of Neurology, 74(1), 20–38. https://doi.org/10.1002/ana.23937

Brown, A.-L., Wilkins, O. G., Keuss, M. J., Hill, S. E., Zanovello, M., Lee, W. C., Bampton, A., Lee, F. C. Y., Masino, L., Qi, Y. A., Bryce-Smith, S., Gatt, A., Hallegger, M., Fagegaltier, D., Phatnani, H., Newcombe, J., Gustavsson, E. K., Seddighi, S., Reyes, J. F., … Fratta, P. (2022). TDP-43 loss and ALS-risk SNPs drive mis-splicing and depletion of UNC13A. Nature, 603(7899), 131–137. https://doi.org/10.1038/s41586-022-04436-3

Cedarbaum, J. M., Stambler, N., Malta, E., Fuller, C., Hilt, D., Thurmond, B., & Nakanishi, A. (1999). The ALSFRS-R: A revised ALS functional rating scale that incorporates assessments of respiratory function. BDNF ALS Study Group (Phase III). Journal of the Neurological Sciences, 169(1–2), 13–21. https://doi.org/10.1016/s0022-510x(99)00210-5

DeJesus-Hernandez, M., Mackenzie, I. R., Boeve, B. F., Boxer, A. L., Baker, M., Rutherford, N. J., Nicholson, A. M., Finch, N. A., Flynn, H., Adamson, J., Kouri, N., Wojtas, A., Sengdy, P., Hsiung, G.-Y. R., Karydas, A., Seeley, W. W., Josephs, K. A., Coppola, G., Geschwind, D. H., … Rademakers, R. (2011). Expanded GGGGCC Hexanucleotide Repeat in Noncoding Region of C9ORF72 Causes Chromosome 9p-Linked FTD and ALS. Neuron, 72(2), 245–256. https://doi.org/10.1016/j.neuron.2011.09.011

Donde, A., Sun, M., Ling, J. P., Braunstein, K. E., Pang, B., Wen, X., Cheng, X., Chen, L., & Wong, P. C. (2019). Splicing repression is a major function of TDP-43 in motor neurons. Acta Neuropathologica, 138(5), 813–826. https://doi.org/10.1007/s00401-019-02042-8

Jeong, Y. H., Ling, J. P., Lin, S. Z., Donde, A. N., Braunstein, K. E., Majounie, E., Traynor, B. J., LaClair, K. D., Lloyd, T. E., & Wong, P. C. (2017). Tdp-43 cryptic exons are highly variable between cell types. Molecular Neurodegeneration, 12(1), 13. https://doi.org/10.1186/s13024-016-0144-x

Josephs, K. A., Murray, M. E., Whitwell, J. L., Parisi, J. E., Petrucelli, L., Jack, C. R., Petersen, R. C., & Dickson, D. W. (2014). Staging TDP-43 pathology in Alzheimer’s disease. Acta Neuropathologica, 127(3), 441–450. https://doi.org/10.1007/s00401-013-1211-9

Josephs, K. A., Whitwell, J. L., Weigand, S. D., Murray, M. E., Tosakulwong, N., Liesinger, A. M., Petrucelli, L., Senjem, M. L., Knopman, D. S., Boeve, B. F., Ivnik, R. J., Smith, G. E., Jack, C. R., Parisi, J. E., Petersen, R. C., & Dickson, D. W. (2014). TDP-43 is a key player in the clinical features associated with Alzheimer’s disease. Acta Neuropathologica, 127(6), 811–824. https://doi.org/10.1007/s00401-014-1269-z

Jumper, J., Evans, R., Pritzel, A., Green, T., Figurnov, M., Ronneberger, O., Tunyasuvunakool, K., Bates, R., Žídek, A., Potapenko, A., Bridgland, A., Meyer, C., Kohl, S. A. A., Ballard, A. J., Cowie, A., Romera-Paredes, B., Nikolov, S., Jain, R., Adler, J., … Hassabis, D. (2021). Highly accurate protein structure prediction with AlphaFold. Nature, 596(7873), 583–589. https://doi.org/10.1038/s41586-021-03819-2

Klim, J. R., Williams, L. A., Limone, F., Juan, I. G. S., Davis-Dusenbery, B. N., Mordes, D. A., Burberry, A., Steinbaugh, M. J., Gamage, K. K., Kirchner, R., Moccia, R., Cassel, S. H., Chen, K., Wainger, B. J., Woolf, C. J., & Eggan, K. (2019). ALS IMPLICATED PROTEIN TDP-43 SUSTAINS LEVELS OF STMN2 A MEDIATOR OF MOTOR NEURON GROWTH AND REPAIR. Nature Neuroscience, 22(2), 167–179. https://doi.org/10.1038/s41593-018-0300-4

Krishnan, G., Raitcheva, D., Bartlett, D., Prudencio, M., McKenna-Yasek, D. M., Douthwright, C., Oskarsson, B. E., Ladha, S., King, O. D., Barmada, S. J., Miller, T. M., Bowser, R., Watts, J. K., Petrucelli, L., Brown, R. H., Kankel, M. W., & Gao, F.-B. (2022). Poly(GR) and poly(GA) in cerebrospinal fluid as potential biomarkers for C9ORF72-ALS/FTD. Nature Communications, 13(1), 2799. https://doi.org/10.1038/s41467-022-30387-4

Lee, E. B., Lee, V. M.-Y., & Trojanowski, J. Q. (2011). Gains or losses: Molecular mechanisms of TDP43-mediated neurodegeneration. Nature Reviews. Neuroscience, 13(1), 38–50. https://doi.org/10.1038/nrn3121

Ling, J. P., Pletnikova, O., Troncoso, J. C., & Wong, P. C. (2015). TDP-43 repression of nonconserved cryptic exons is compromised in ALS-FTD. Science (New York, N.Y.), 349(6248), 650–655. https://doi.org/10.1126/science.aab0983

Ling, J. P., Wilks, C., Charles, R., Leavey, P. J., Ghosh, D., Jiang, L., Santiago, C. P., Pang, B., Venkataraman, A., Clark, B. S., Nellore, A., Langmead, B., & Blackshaw, S. (2020). ASCOT identifies key regulators of neuronal subtype-specific splicing. Nature Communications, 11(1), Article 1. https://doi.org/10.1038/s41467-019-14020-5

Ma, X. R., Prudencio, M., Koike, Y., Vatsavayai, S. C., Kim, G., Harbinski, F., Briner, A., Rodriguez, C. M., Guo, C., Akiyama, T., Schmidt, H. B., Cummings, B. B., Wyatt, D. W., Kurylo, K., Miller, G., Mekhoubad, S., Sallee, N., Mekonnen, G., Ganser, L., … Gitler, A. D. (2022). TDP-43 represses cryptic exon inclusion in the FTD–ALS gene UNC13A. Nature, 603(7899), 124–130. https://doi.org/10.1038/s41586-022-04424-7

Masaki, K., Sonobe, Y., Ghadge, G., Pytel, P., Lépine, P., Pernin, F., Cui, Q.-L., Antel, J. P., Zandee, S., Prat, A., & Roos, R. P. (2020). RNA-binding protein altered expression and mislocalization in MS. Neurology(R) Neuroimmunology & Neuroinflammation, 7(3), e704. https://doi.org/10.1212/NXI.0000000000000704

McKee, A. C., Gavett, B. E., Stern, R. A., Nowinski, C. J., Cantu, R. C., Kowall, N. W., Perl, D. P., Hedley-Whyte, E. T., Price, B., Sullivan, C., Morin, P., Lee, H.-S., Kubilus, C. A., Daneshvar, D. H., Wulff, M., & Budson, A. E. (2010). TDP-43 proteinopathy and motor neuron disease in chronic traumatic encephalopathy. Journal of Neuropathology and Experimental Neurology, 69(9), 918–929. https://doi.org/10.1097/NEN.0b013e3181ee7d85

Melamed, Z., Lopez-Erauskin, J., Baughn, M. W., Zhang, O., Drenner, K., Sun, Y., Freyermuth, F., McMahon, M. A., Beccari, M. S., Artates, J., Ohkubo, T., Rodriguez, M., Lin, N., Wu, D., Bennett, C. F., Rigo, F., Da Cruz, S., Ravits, J., Lagier-Tourenne, C., & Cleveland, D. W. (2019). Premature polyadenylation-mediated loss of stathmin-2 is a hallmark of TDP-43-dependent neurodegeneration. Nature Neuroscience, 22(2), 180–190. https://doi.org/10.1038/s41593-018-0293-z

Meneses, A., Koga, S., O’Leary, J., Dickson, D. W., Bu, G., & Zhao, N. (2021). TDP-43 Pathology in Alzheimer’s Disease. Molecular Neurodegeneration, 16(1), 84. https://doi.org/10.1186/s13024-021-00503-x

Mitsumoto, H., Brooks, B. R., & Silani, V. (2014). Clinical trials in amyotrophic lateral sclerosis: Why so many negative trials and how can trials be improved? The Lancet Neurology, 13(11), 1127–1138. https://doi.org/10.1016/S1474-4422(14)70129-2

Nelson, P. T., Brayne, C., Flanagan, M. E., Abner, E. L., Agrawal, S., Attems, J., Castellani, R. J., Corrada, M. M., Cykowski, M. D., Di, J., Dickson, D. W., Dugger, B. N., Ervin, J. F., Fleming, J., Graff-Radford, J., Grinberg, L. T., Hokkanen, S. R. K., Hunter, S., Kapasi, A., … Schneider, J. A. (2022). Frequency of LATE neuropathologic change across the spectrum of Alzheimer’s disease neuropathology: Combined data from 13 community-based or population-based autopsy cohorts. Acta Neuropathologica, 144(1), 27–44. https://doi.org/10.1007/s00401-022-02444-1

Nelson, P. T., Dickson, D. W., Trojanowski, J. Q., Jack, C. R., Boyle, P. A., Arfanakis, K., Rademakers, R., Alafuzoff, I., Attems, J., Brayne, C., Coyle-Gilchrist, I. T. S., Chui, H. C., Fardo, D. W., Flanagan, M. E., Halliday, G., Hokkanen, S. R. K., Hunter, S., Jicha, G. A., Katsumata, Y., … Schneider, J. A. (2019). Limbic-predominant age-related TDP-43 encephalopathy (LATE): Consensus working group report. Brain: A Journal of Neurology, 142(6), 1503–1527. https://doi.org/10.1093/brain/awz099

Neumann, M., Sampathu, D. M., Kwong, L. K., Truax, A. C., Micsenyi, M. C., Chou, T. T., Bruce, J., Schuck, T., Grossman, M., Clark, C. M., McCluskey, L. F., Miller, B. L., Masliah, E., Mackenzie, I. R., Feldman, H., Feiden, W., Kretzschmar, H. A., Trojanowski, J. Q., & Lee, V. M.-Y. (2006). Ubiquitinated TDP-43 in Frontotemporal Lobar Degeneration and Amyotrophic Lateral Sclerosis. Science, 314(5796), 130–133.

Offit, M. B., Wu, T., Floeter, M. K., & Lehky, T. J. (2020). Electrical impedance myography (EIM) in a natural history study of C9ORF72 mutation carriers. Amyotrophic Lateral Sclerosis & Frontotemporal Degeneration, 21(5–6), 445–451. https://doi.org/10.1080/21678421.2020.1752247

Prudencio, M., Humphrey, J., Pickles, S., Brown, A.-L., Hill, S. E., Kachergus, J. M., Shi, J., Heckman, M. G., Spiegel, M. R., Cook, C., Song, Y., Yue, M., Daughrity, L. M., Carlomagno, Y., Jansen-West, K., de Castro, C. F., DeTure, M., Koga, S., Wang, Y.-C., … Petrucelli, L. (2020). Truncated stathmin-2 is a marker of TDP-43 pathology in frontotemporal dementia. The Journal of Clinical Investigation, 130(11), 6080–6092. https://doi.org/10.1172/JCI139741

Renton, A. E., Majounie, E., Waite, A., Simón-Sánchez, J., Rollinson, S., Gibbs, J. R., Schymick, J. C., Laaksovirta, H., van Swieten, J. C., Myllykangas, L., Kalimo, H., Paetau, A., Abramzon, Y., Remes, A. M., Kaganovich, A., Scholz, S. W., Duckworth, J., Ding, J., Harmer, D. W., … Traynor, B. J. (2011). A hexanucleotide repeat expansion in C9ORF72 is the cause of chromosome 9p21-linked ALS-FTD. Neuron, 72(2), 257–268. https://doi.org/10.1016/j.neuron.2011.09.010

Robinson, J. L., Lee, E. B., Xie, S. X., Rennert, L., Suh, E., Bredenberg, C., Caswell, C., Van Deerlin, V. M., Yan, N., Yousef, A., Hurtig, H. I., Siderowf, A., Grossman, M., McMillan, C. T., Miller, B., Duda, J. E., Irwin, D. J., Wolk, D., Elman, L., … Trojanowski, J. Q. (2018). Neurodegenerative disease concomitant proteinopathies are prevalent, age-related and APOE4-associated. Brain: A Journal of Neurology, 141(7), 2181–2193. https://doi.org/10.1093/brain/awy146

Tan, Q., Yalamanchili, H. K., Park, J., De Maio, A., Lu, H.-C., Wan, Y.-W., White, J. J., Bondar, V. V., Sayegh, L. S., Liu, X., Gao, Y., Sillitoe, R. V., Orr, H. T., Liu, Z., & Zoghbi, H. Y. (2016). Extensive cryptic splicing upon loss of RBM17 and TDP43 in neurodegeneration models. Human Molecular Genetics, 25(23), 5083–5093. https://doi.org/10.1093/hmg/ddw337

Taylor, J. P., Brown, R. H., & Cleveland, D. W. (2016). Decoding ALS: From genes to mechanism. Nature, 539(7628), 197–206. https://doi.org/10.1038/nature20413

Traynor, B. J., Codd, M. B., Corr, B., Forde, C., Frost, E., & Hardiman, O. (2000). Amyotrophic lateral sclerosis mimic syndromes: A population-based study. Archives of Neurology, 57(1), 109–113. https://doi.org/10.1001/archneur.57.1.109

Vatsavayai, S. C., Yoon, S. J., Gardner, R. C., Gendron, T. F., Vargas, J. N. S., Trujillo, A., Pribadi, M., Phillips, J. J., Gaus, S. E., Hixson, J. D., Garcia, P. A., Rabinovici, G. D., Coppola, G., Geschwind, D. H., Petrucelli, L., Miller, B. L., & Seeley, W. W. (2016). Timing and significance of pathological features in *C9orf72* expansion-associated frontotemporal dementia. Brain, 139(12), 3202–3216. https://doi.org/10.1093/brain/aww250

Venkataraman, A., Yang, K., Irizarry, J., Mackiewicz, M., Mita, P., Kuang, Z., Xue, L., Ghosh, D., Liu, S., Ramos, P., Hu, S., Bayron Kain, D., Keegan, S., Saul, R., Colantonio, S., Zhang, H., Behn, F. P., Song, G., Albino, E., … Blackshaw, S. (2018). A toolbox of immunoprecipitation-grade monoclonal antibodies to human transcription factors. Nature Methods, 15(5), Article 5. https://doi.org/10.1038/nmeth.4632

Zhang, Y.-J., Xu, Y.-F., Cook, C., Gendron, T. F., Roettges, P., Link, C. D., Lin, W.-L., Tong, J., Castanedes-Casey, M., Ash, P., Gass, J., Rangachari, V., Buratti, E., Baralle, F., Golde, T. E., Dickson, D. W., & Petrucelli, L. (2009). Aberrant cleavage of TDP-43 enhances aggregation and cellular toxicity. Proceedings of the National Academy of Sciences, 106(18), 7607–7612. https://doi.org/10.1073/pnas.0900688106

